## Supplemental Information 1 for "Odor-evoked transcriptomics of *Aedes aegypti* mosquitoes"

### Supporting Information

#### KEY RESOURCES TABLE

| Reagents and Resources | Source | Identifiers |
|---|---|---|
| Tetramin tropical fish food | Tetra | Catalog#16152 |
| BugDorm Rearing cage | BioQuip | Catalog#1452 |
| Drosophila food vials | GeneseeScientific | Catalog#32-117BF |
| Cottonballs | ThermoFisherScientific | Catalog#22-456-883 |
| polytubing | Uline | ModelNo.S-3521 |
| Ant habitat | KristalEduacational | Item#6130 |
| Disposable pellet pestles | FisherScientific | Catalog#12-141-364 |
| **Chemicals** | | |
| Guanidinethiocyanate | Sigma-Aldrich | C.A.S.593-84-0 |
| Chloroform | FisherScientific | C.A.S.67-66-3 |
| Sarkosyl | FisherScientific | C.A.S.137-16-6 |
| 2-mercapthoethanol | Sigma-Aldrich | C.A.S:60-24-2 |
| 1-octen-3-ol | Sigma-Aldrich | C.A.S.3391-86-4 |
| Mineraloil | ThermoFisherScientific | Catalog#O122-1 |
| **Reagents** | | |
| RNAlaterstabilizationsolution | Invitrogen | Catalog#AM7020 |
| RNAidKit | MPBio | catalog#111007200 |
| AmplitaqGold360PCRmastermix | ThermoFisherScientific | Catalog#4398881 |
| TurboDNA-free™removalKit | Invitrogen | Catalog#AM1906 |
| usingVersocDNASynthesisKit | ThermoFisherScientific | CatalogAB1453A |
| RNaseCocktailEnzymeMix | ThermoFisherScientific | Catalog#AM2286 |
| TaqMan2Xuniversalmastermix | Thermo-FisherScientific | Catalog#4324018 |
| CustomTaqManprobes | ThermoFisherScientific | Catalog#4331348 |
| **Software** | | |
| ThermoScientificNanoDrop2000c | ThermoFisherScientific | v1.6.198 |
| RealtimePCRsystem | AppliedBiosystems | 7500 |
| SDSsoftware | AppliedBiosystems | v1.4.1 |
| FASTQC | https://www.bioinformatics.babraham.ac.uk/projects/fastqc/ | v0.11.5 |
| Trimmomatic | sadellab.org | v0.36 |
| Vectorbase | vectorbase.com | release55 |

| | | |
|---|---|---|
| tximport | Bioconductor | v1.18.0 |
| DESeq2 | Bioconductor | v1.34.0 |
| RStudio | https://www.rstudio.com/ | v1.2.1335 |
| Salmon | https://combine-lab.github.io/salmon/ | v9.1 |
| GraphPadPrism | GraphPad | v8 |
| ComplexHeatmap | Bioconductor | 2.13.1 |
| ggplot2 | Tidyverse | 3.4.0 |
| Mol3DViewer | https://www.rcsb.org/3d-view | 3.4.1 |
| LGAprogram | http://proteinmodel.org/AS2TS/LGA/lga.html | License_39633_220526_00t |

| Time | Concentration | Alive | Moribund | Dead |
|---|---|---|---|---|
| 6 hour | Mineral oil | 5-5-5 | 0 | 0 |
| 6 hour | 4% 1-octen-3-ol | 5-5-5 | 0 | 0 |
| 6 hour | 2% 1-octen-3-ol | 5-5-5 | 0 | 0 |
| 6 hour | 1% 1-octen-3-ol | 5-5-5 | 0 | 0 |
| 6 hour | 0.5% 1-octen-3-ol | 5-5-5 | 0 | 0 |
| 24 hour | Mineral oil | 5-5-5 | 0 | 0 |
| 24 hour | 4% 1-octen-3-ol | 0 | 0 | 5-5-5 |
| 24 hour | 2% 1-octen-3-ol | 0 | 5-5-5 | 0 |
| 24 hour | 1%1-octen-3-ol | 5-5-5 | 0 | 0 |
| 24 hour | 0.5% 1-octen-3-ol | 5-5-5 | 0 | 0 |

S1 Table A. no-observed-adverse-effect level (NOAEL) assay for 6 and 24 hour 1-octen-3-ol exposure.

| **OR8 probe** |
|---|
| Forward Primer Sequence: GCGATGAAATTCTCCGTAGTTTCG |
| Reverse Primer Sequence: GCAAGGTGAAGTATGAAAACGACGTA |
| Reporter 1 Sequence: AAGATCGCCGAATAGC |
| **RPL32_Probe: (housekeeping gene)** |
| Forward Primer Sequence: ATCAGTCCGATCGCTATGACAAG |
| Reverse Primer Sequence: GGTTGTCAATACCTTTCGGCTTAC |
| Reporter 1 Sequence: TTGCCCCCAACTGGC |

S1 Table B. Primer Probes used in RT-qPCR evaluation of Or8 mRNA.

| | 1hr | 3hr | 6hr | 12hr | 24hr |
|---|---|---|---|---|---|
| 1 | 1.0148 | 0.7599 | 0.2347 | 0.1878 | 0.0279 |
| 2 | 0.9915 | 0.5764 | 0.2348 | 0.1826 | 0.0601 |
| 3 | 0.9598 | 0.3950 | 0.1949 | 0.1575 | 0.0381 |
| 4 | 1.1372 | 0.8068 | 0.1855 | 0.1573 | 0.0795 |
| 5 | 1.0469 | 0.8606 | 0.2118 | 0.1559 | 0.0670 |
| 6 | 1.1508 | 0.6365 | 0.2932 | 0.1449 | 0.0790 |
| 7 | 1.0231 | 0.8030 | 0.0907 | 0.2400 | 0.3299 |
| 8 | 0.9826 | 1.0955 | 0.1346 | 0.2528 | 0.2809 |
| 9 | 0.9319 | 0.6303 | 0.0955 | 0.2977 | 0.1569 |

S1 Table C. Relative fold change values of Or8 mRNA when exposed to 1% 1-octen-3-ol at varying lengths of time.

| | 1 | $10^{-1}$ | $10^{-2}$ | $10^{-3}$ | $10^{-4}$ | $10^{-5}$ | $10^{-6}$ | $10^{-7}$ | $10^{-8}$ |
|---|---|---|---|---|---|---|---|---|---|
| 1 | 0.1542 | 0.2846 | 0.3146 | 0.3496 | 0.5019 | 0.2550 | 0.7830 | 0.7572 | 1.2710 |
| 2 | 0.1818 | 0.2469 | 0.3348 | 0.4253 | 0.3460 | 0.2246 | 0.5770 | 0.5825 | 1.4288 |
| 3 | 0.1227 | 0.2276 | 0.3376 | 0.3013 | 0.4737 | 0.2810 | 0.7312 | 0.5926 | 1.1848 |
| 4 | 0.1696 | 0.3718 | 0.1768 | 0.2872 | 0.4715 | 0.4008 | 0.4360 | 0.4604 | 0.9825 |
| 5 | 0.1704 | 0.3851 | 0.1384 | 0.2176 | 0.6290 | 0.3093 | 0.5725 | 0.4827 | 0.9071 |
| 6 | 0.1439 | 0.4212 | 0.1227 | 0.2410 | 0.4485 | 0.3245 | 0.5845 | 0.5161 | 0.9154 |
| 7 | 0.1568 | 0.3980 | 0.4692 | 0.2159 | 0.4958 | 0.9036 | 1.0047 | 0.8602 | 0.5484 |
| 8 | 0.1176 | 0.4984 | 0.3178 | 0.1857 | 0.5634 | 0.5619 | 0.9295 | 0.9779 | 0.5540 |
| 9 | 0.1343 | 0.2964 | 0.3509 | 0.2218 | 0.3984 | 0.6536 | 0.9610 | 1.0775 | 0.4543 |
| 10 | | | | | | 0.2540 | | 0.9227 | |
| 11 | | | | | | 0.3978 | | 0.9485 | |
| 12 | | | | | | 0.3247 | | 1.0484 | |

S1 Table D. Relative fold change values of Or8 mRNA levels to varying concentrations of 1-octen-3-ol after 6 hours.

| | 6 hour exposed | 6 hour exposed + 6 hour recovery |
|---|---|---|
| 1 | 0.1973 | 0.3056 |
| 2 | 0.1667 | 0.3881 |
| 3 | 0.2009 | 0.3152 |
| 4 | 0.1616 | 0.6633 |
| 5 | 0.1638 | 0.7421 |
| 6 | 0.1722 | 0.6296 |
| 7 | 0.3429 | 0.9598 |
| 8 | 0.3396 | 1.1152 |
| 9 | 0.3756 | 1.0557 |

S1 Table E. Relative fold change values of Or8 mRNA when exposed to 1% 1-octen-3-ol for 6 hours; then given an additional 6 hour to recover.

| | Control | 1% 1-octen-3-ol |
|---|---|---|
| 1 | 0.9204 | 0.4977 |
| 2 | 1.0730 | 0.4210 |
| 3 | 0.9888 | 0.4973 |
| 4 | 1.2064 | 0.5203 |
| 5 | 1.4890 | 0.5231 |
| 6 | 1.3571 | 0.5801 |
| 7 | 0.7086 | 0.2798 |
| 8 | 0.8969 | 0.1531 |
| 9 | 0.6610 | 0.2987 |

S1 Table F. Large assay validation of Or8 mRNA response to 1-octen-3-ol exposure

| SampleIDs | Tissuetype | ObservedFragments | UnMappedReads | CountedReads | MappingRate |
|---|---|---|---|---|---|
| OctT1 | Antenna,Maxillary,Proboscis | 277,118,382 | 64,657,152 | 212,461,230 | 76.67% |
| OctT2 | Antenna,Maxillary,Proboscis | 265,657,872 | 86,994,300 | 178,663,572 | 67.25% |
| OctT3 | Antenna,Maxillary,Proboscis | 266,863,157 | 72,489,769 | 194,373,388 | 72.84% |
| OctC1 | Antenna,Maxillary,Proboscis | 206,372,686 | 42,595,443 | 163,777,243 | 79.36% |
| OctC2 | Antenna,Maxillary,Proboscis | 291,393,839 | 58,927,013 | 232,466,826 | 79.78% |
| OctC3 | Antenna,Maxillary,Proboscis | 236,756,732 | 58,216,704 | 178,540,028 | 75.41% |
| Totals | | 1,544,162,668 | 383,880,381 | 1,160,282,287 | 75.22% |

S1 Table G. Summary of RNA-seq samples and their mapping rate

| Conditions | GeneRatio | p.adjust | Counts | Ontology |
|---|---|---|---|---|
| Integral component of membrane | 0.073384447 | 2.06E-06 | 201 | CC |
| Intrinsic component of membrane | 0.073250729 | 2.06E-06 | 201 | CC |
| Membrane | 0.0687251 | 0.000121933 | 207 | CC |
| Sensory perception of smell | 0.259259259 | 2.30E-07 | 21 | BP |
| Detection of chemical stimulus | 0.225 | 1.68E-05 | 18 | BP |
| Detection of stimulus | 0.195652174 | 0.000110272 | 18 | BP |
| System process | 0.149350649 | 0.000330205 | 23 | BP |
| Response to chemical | 0.122340426 | 0.00773979 | 23 | BP |
| Serine family amino acid metabolic process | 0.454545455 | 0.007790979 | 5 | BP |
| Odorant binding | 0.292134831 | 7.62E-23 | 52 | MF |
| Iron ion binding | 0.204878049 | 2.16E-12 | 42 | MF |
| Monooxygenase activity | 0.226993865 | 2.16E-12 | 37 | MF |
| Oxidoreductase activity, acting on paired donors | 0.201030928 | 2.10E-11 | 39 | MF |
| Heme binding | 0.190721649 | 3.68E-10 | 37 | MF |
| Tetrapyrrole binding | 0.18974359 | 3.68E-10 | 37 | MF |
| Oxidoreductase activity | 0.112107623 | 1.01E-08 | 75 | MF |
| Olfactory receptor activity | 0.256410256 | 9.79E-08 | 20 | MF |
| Serine hydrolase activity | 0.096774194 | 0.009564437 | 36 | MF |
| Serine-type peptidase activity | 0.096774194 | 0.009564437 | 36 | MF |
| Drug metabolism - cytochromeP450 | 0.166666667 | 1.07E-06 | 37 | KEGG |
| Retinol metabolism | 0.151750973 | 3.03E-06 | 39 | KEGG |
| Metabolism of xenobiotics by cytochrome P450 | 0.171270718 | 3.85E-06 | 31 | KEGG |
| Steroid hormone biosynthesis | 0.169590643 | 1.01E-05 | 29 | KEGG |
| Caffeine metabolism | 0.225 | 2.75E-05 | 18 | KEGG |
| Indole alkaloid biosynthesis | 0.172932331 | 9.42E-05 | 23 | KEGG |
| Fatty acid degradation | 0.149068323 | 0.000689445 | 24 | KEGG |
| Ascorbate and aldarate metabolism | 0.128099174 | 0.00088728 | 31 | KEGG |
| Styrene degradation | 0.185185185 | 0.001502926 | 15 | KEGG |
| Isoflavonoid biosynthesis | 0.191176471 | 0.002879573 | 13 | KEGG |
| Brassinosteroid biosynthesis | 0.172839506 | 0.004529185 | 14 | KEGG |

S1 Table H. Summary of Gene Ontology (GO) and KEGG pathways analysis of DE genes.

| | AaOr8 | TaOr8 | AgOr8 | CqOr114 | CqOr118 | AgOr4 | AgOr20 | CqOr1 | AgOr59 | AalOr10 | AalOr88 | BdorOr13a | BminOr3 | DmelOr13a | DmelOr85c |
|---|---|---|---|---|---|---|---|---|---|---|---|---|---|---|---|
| **AaOr4** | 87.74 | 87.05 | 86.61 | 81.38 | 83.38 | 56.47 | 57.98 | 60.89 | 56.43 | 75.51 | 52.83 | 80.15 | 82.66 | 83.07 | 84.15 |
| **AaOr8** | 100 | 85.38 | 85.48 | 75.57 | 80.16 | 48.80 | 54.13 | 56.48 | 49.55 | 59.15 | 45.87 | 74.42 | 74.64 | 74.72 | 77.31 |
| **AaOr10** | 72.98 | 71.89 | 69.97 | 63.37 | 67.58 | 62.66 | 61.66 | 66.47 | 60.12 | 99.44 | 51.32 | 72.47 | 72.61 | 71.64 | 71.28 |
| **AaOr11** | 64.21 | 65.30 | 65.82 | 59.09 | 59.67 | 51.02 | 56.35 | 60.04 | 57.85 | 66.29 | 51.33 | 63.39 | 64.40 | 62.82 | 63.54 |
| **AaOr13** | 54.35 | 63.32 | 60.65 | 60.91 | 59.44 | 86.43 | 87.26 | 89.13 | 54.06 | 69.35 | 53.01 | 54.32 | 53.47 | 59.25 | 62.85 |
| **AaOr28** | 80.76 | 93.34 | 92.00 | 90.07 | 87.09 | 55.04 | 60.78 | 65.57 | 54.22 | 72.72 | 51.39 | 76.18 | 74.59 | 80.56 | 87.46 |
| **AaOr29** | 36.25 | 37.27 | 41.13 | 63.07 | 39.71 | 41.18 | 37.38 | 40.48 | 40.41 | 35.58 | 36.87 | 35.59 | 47.27 | 65.50 | 62.86 |
| **AaOr31** | 68.67 | 78.86 | 78.19 | 60.32 | 73.77 | 55.07 | 56.18 | 62.63 | 53.70 | 84.60 | 51.23 | 67.32 | 65.78 | 69.14 | 67.97 |
| **AaOr36** | 51.19 | 71.38 | 58.20 | 56.54 | 55.85 | 56.94 | 51.68 | 57.27 | 55.61 | 50.85 | 61.77 | 48.79 | 62.99 | 60.81 | 56.65 |
| **AaOr51** | 23.95 | 27.77 | 27.04 | 31.93 | 31.53 | 36.94 | 33.25 | 34.91 | 31.78 | 30.02 | 28.88 | 26.45 | 25.77 | 28.01 | 27.82 |
| **AaOr52** | 50.86 | 59.71 | 58.70 | 57.70 | 55.86 | 57.43 | 58.92 | 61.02 | 62.38 | 63.93 | 72.44 | 54.41 | 53.69 | 56.21 | 61.46 |
| **AaOr66** | 52.84 | 61.52 | 59.28 | 60.19 | 57.36 | 64.79 | 74.83 | 78.84 | 50.48 | 64.98 | 50.05 | 50.74 | 50.38 | 57.35 | 65.06 |
| **AaOr69** | 53.40 | 62.76 | 57.86 | 60.24 | 57.33 | 73.08 | 72.70 | 76.59 | 51.33 | 67.90 | 50.45 | 49.05 | 48.55 | 57.42 | 61.81 |
| **AaOr70** | 56.38 | 63.65 | 62.94 | 67.69 | 62.13 | 64.84 | 65.25 | 76.42 | 53.68 | 64.16 | 50.19 | 48.85 | 49.60 | 57.60 | 63.61 |
| **AaOr72** | 44.84 | 54.73 | 53.93 | 55.97 | 54.02 | 53.82 | 54.77 | 58.55 | 65.25 | 59.92 | 58.65 | 48.83 | 48.52 | 49.17 | 53.78 |
| **AaOr85** | 51.58 | 60.29 | 59.61 | 55.11 | 55.08 | 57.12 | 56.02 | 58.05 | 61.30 | 63.50 | 65.05 | 49.24 | 46.79 | 54.28 | 55.24 |
| **AaOr88** | 44.86 | 52.41 | 51.46 | 50.26 | 48.60 | 53.41 | 56.69 | 56.02 | 57.00 | 52.60 | 96.13 | 45.25 | 44.45 | 48.86 | 50.77 |
| **AaOr94** | 41.47 | 49.23 | 47.45 | 49.43 | 49.27 | 53.89 | 49.68 | 53.17 | 57.02 | 52.57 | 64.06 | 42.14 | 39.14 | 43.98 | 47.56 |
| **AaOr100** | 46.05 | 55.10 | 52.84 | 52.56 | 50.36 | 49.98 | 55.20 | 54.55 | 58.32 | 55.21 | 82.61 | 46.01 | 44.98 | 53.25 | 53.98 |
| **AaOr107** | 42.65 | 50.47 | 48.15 | 54.58 | 51.95 | 50.08 | 52.04 | 53.36 | 49.65 | 52.64 | 59.50 | 40.88 | 40.21 | 45.01 | 54.16 |
| **AaOr112** | 40.97 | 47.83 | 45.81 | 49.74 | 46.24 | 44.22 | 46.74 | 46.67 | 48.79 | 49.75 | 59.84 | 39.62 | 38.43 | 42.55 | 49.27 |
| **AaOr114** | 39.41 | 46.38 | 44.90 | 45.26 | 44.23 | 49.11 | 47.99 | 51.50 | 52.40 | 49.05 | 58.13 | 39.98 | 39.11 | 44.00 | 48.48 |
| **AaOr115** | 47.52 | 57.07 | 54.55 | 52.97 | 51.16 | 47.83 | 53.77 | 55.44 | 56.99 | 54.69 | 78.26 | 48.51 | 45.96 | 50.72 | 55.84 |

S1 Table I. structure similarity scores (LGA_S) of Alphafold2 generated candidate receptors superimposed onto known dipteran 1-octen-3-ol receptors.
